## Supplementary figures and images for "Sirtuin-1 Sensitive Lysine-136 Acetylation Drives Phase Separation and Pathological Aggregation of TDP-43"

### Supplementary Figure 1

Supplementary figure 1

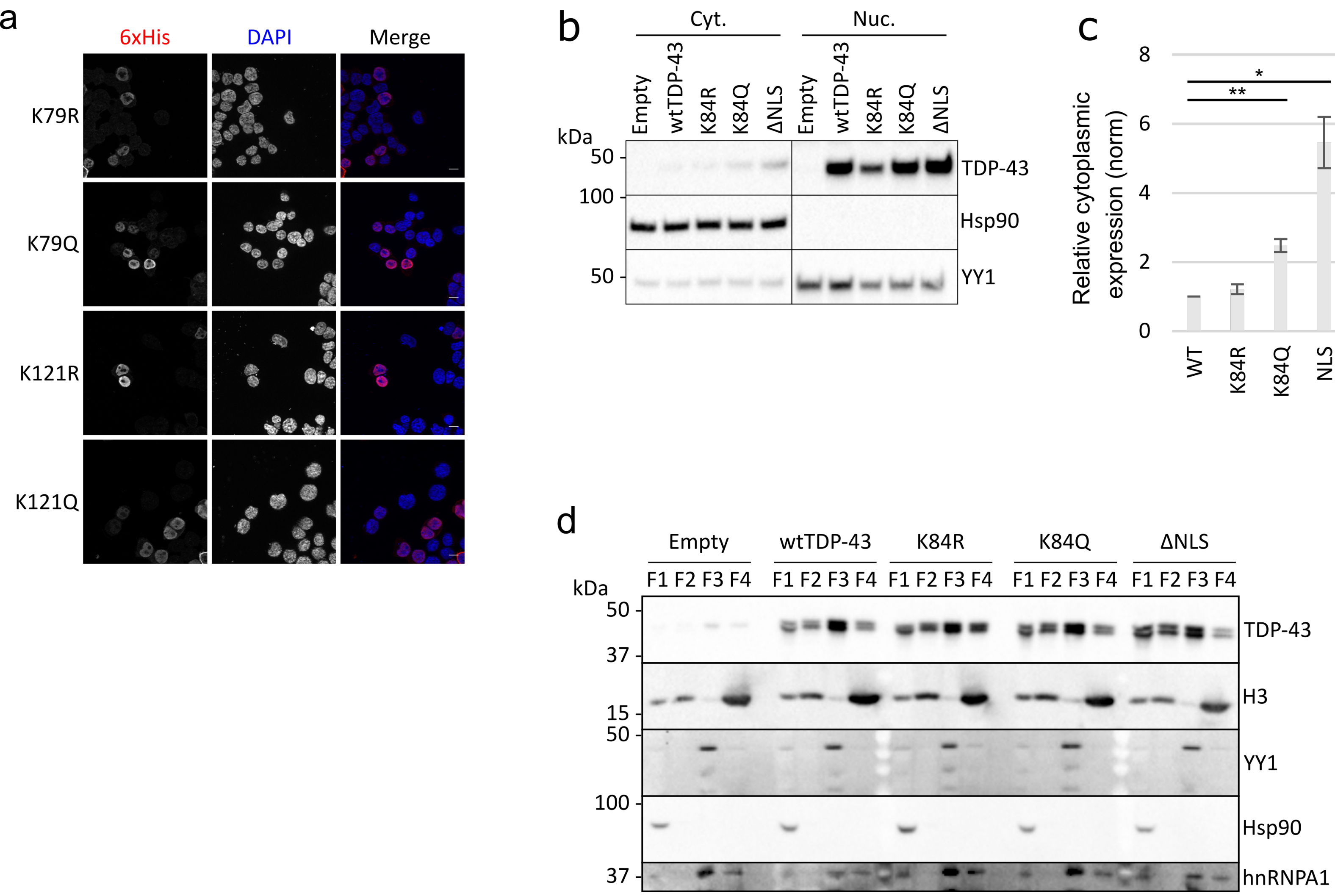

### Supplementary figure 2

#### Supplementary figure 2

a

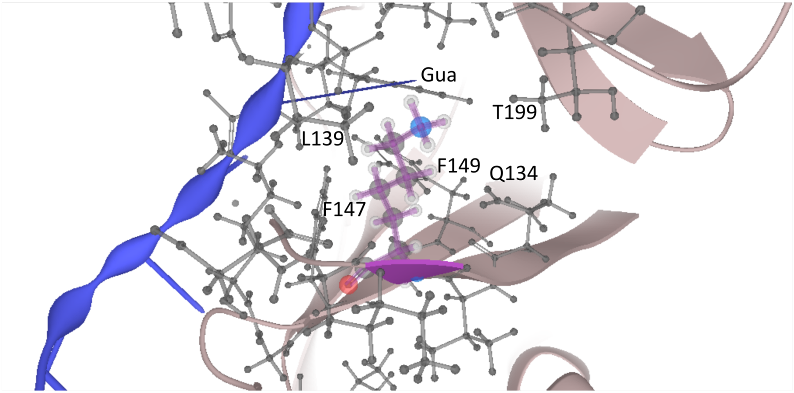


b

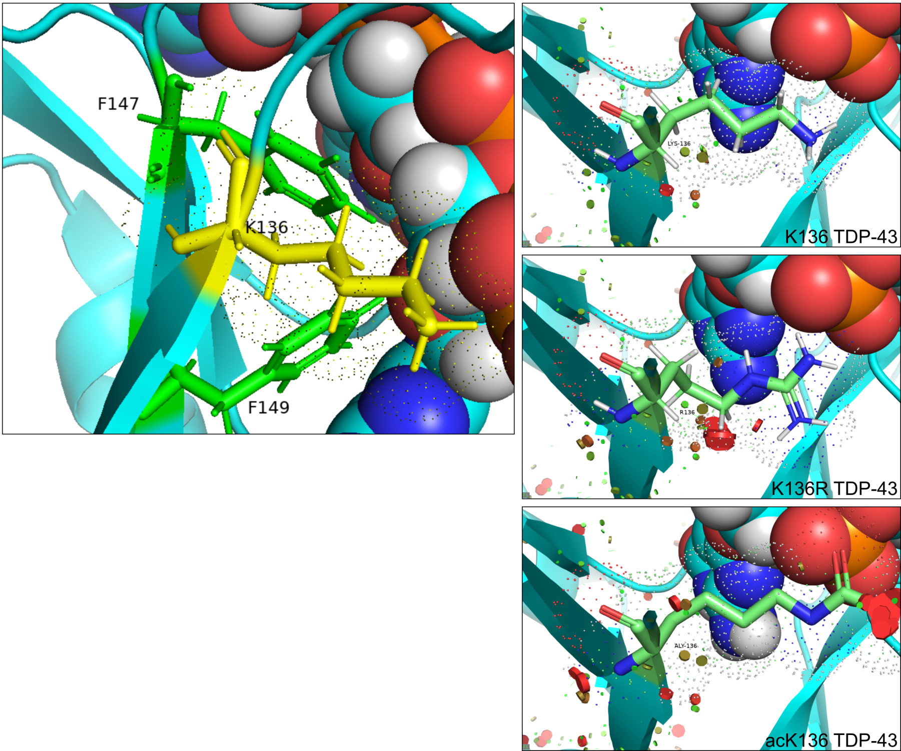

### Supplementary figure 3

Supplementary figure 3

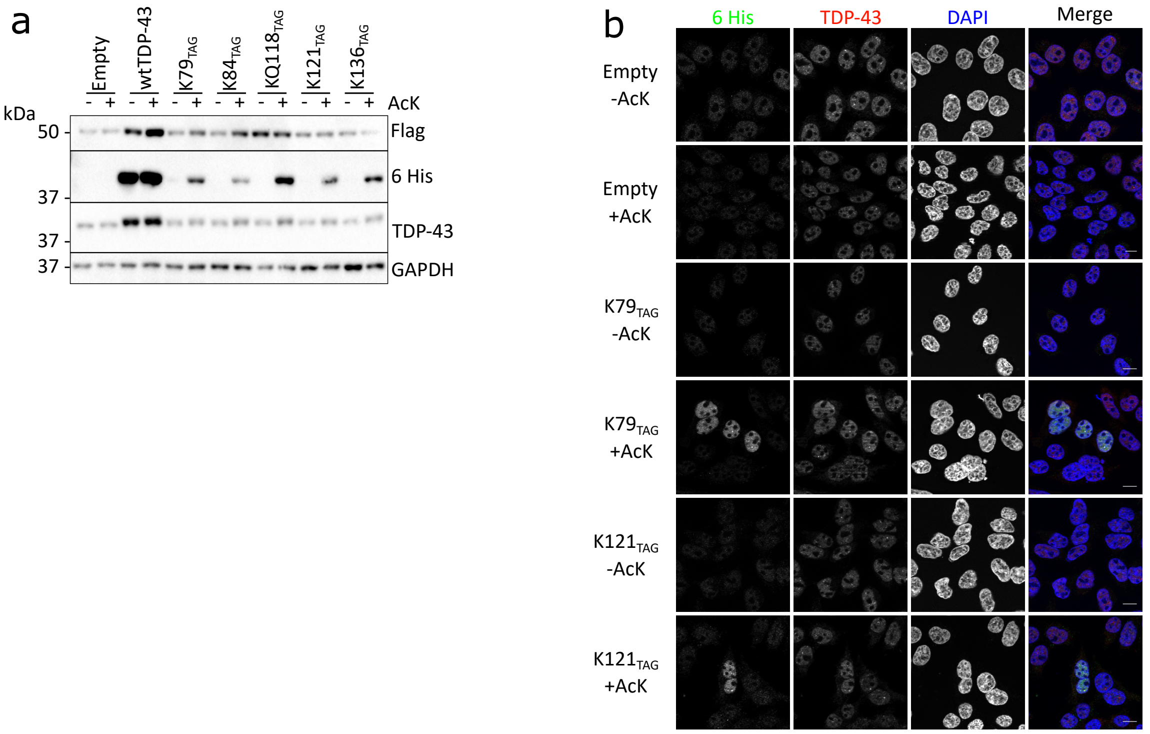

### Supplementary figure 4

Supplementary figure 4

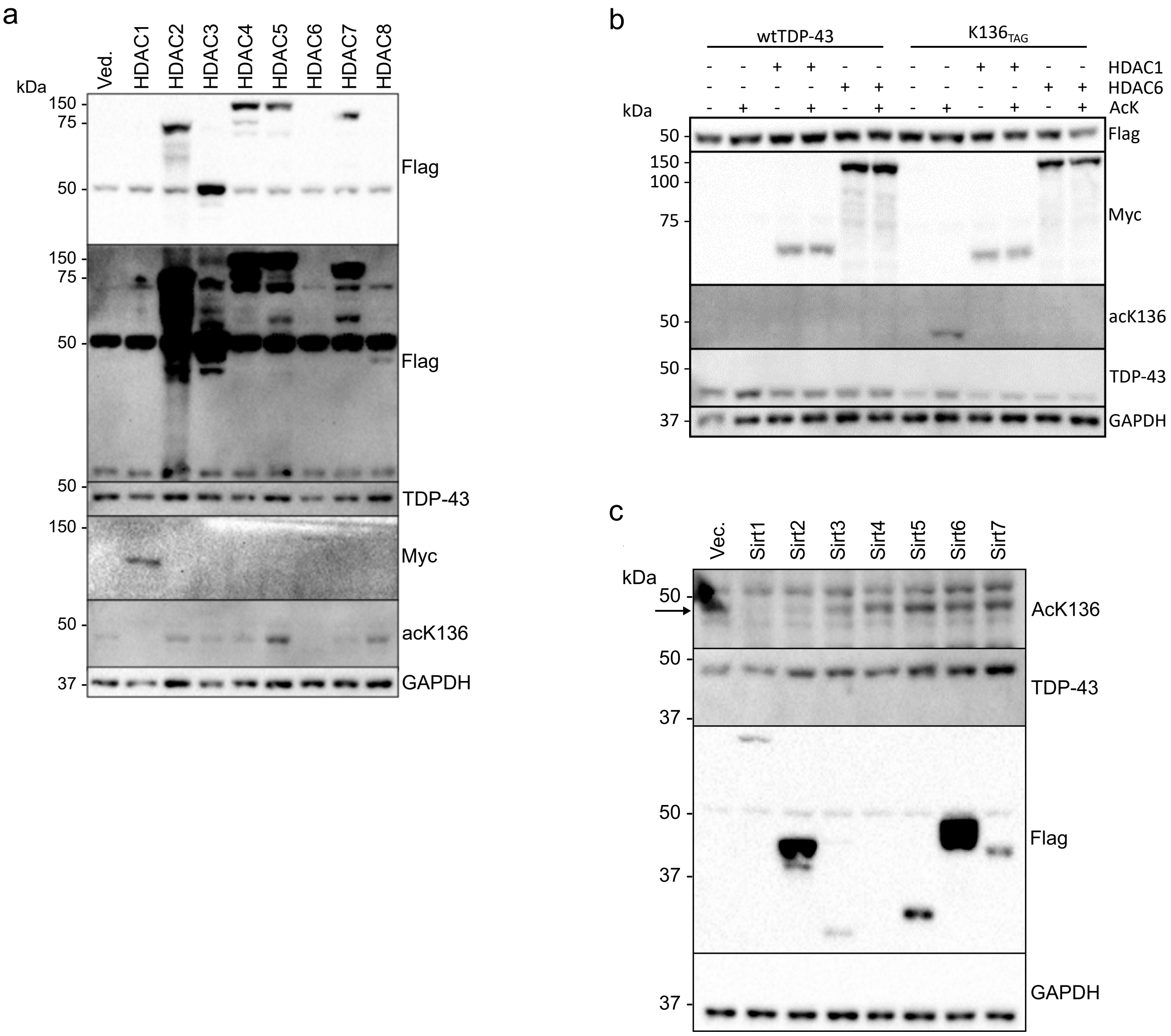
