## Supplementary table 1 for "Sirtuin-1 Sensitive Lysine-136 Acetylation Drives Phase Separation and Pathological Aggregation of TDP-43"

| Peptide sequence | Peptide lysine | TDP-43 residue | Mascot identity score |
|---|---|---|---|
| RLVEGILHAPDAGWGNLVYVVNYP<u>K</u>DNKR | K25 | K79 | 27.2 |
| R<u>K</u>MDETDASSAVKV | K2 | K84 | 28.3 |
| RAVQ<u>K</u>TSDLIVLGLPWKT | K5 | K121 | 25.0 |
| KEYFSTFGEVLMVQV<u>K</u>KD | K17 | K136 | 30.0 |
