## Supplementary table 2 for "Sirtuin-1 Sensitive Lysine-136 Acetylation Drives Phase Separation and Pathological Aggregation of TDP-43"

| Primer | Sequence |
| --- | --- |
| TDPΔGRD NotI reverse | ccccgcggccgcctaacttctttctaactgtctattgctattg |
| TDP-43 6His BamHI reverse | cagcggccgcggatccttaatggtgatggtgatgatgcattccccagccagaag |
| TDP-43 6His NheI forward | gctagccatcatcaccatcaccatatgtctgaatatattc |
| TDP-43 BamHI forward | ggggggatccgatgtctgaatatattcgggtaacc |
| TDP-43 Bsp120 forward | gggggggcccaccatgtctgaatatattcgggtaaccg |
| TDP-43 HindIII reverse | ccccaagcttctacattccccagccagaag |
| TDP43 K121Q forward | ccgaacaggacctgcaagagtattttagtac |
| TDP43 K121Q reverse | gtactaaaatactcttgcaggtcctgttcgg |
| TDP43 K121R forward | ccgaacaggacctgagagagtattttagtac |
| TDP43 K121R reverse | gtactaaaatactctctcaggtcctgttcgg |
| TDP43 K136Q forward | ggtgcaggtccagaaagatcttaagactgg |
| TDP43 K136Q reverse | ccagtcttaagatctttctggacctgcacc |
| TDP43 K136R forward | ggtgcaggtcaggaaagatcttaagactgg |
| TDP43 K136R reverse | ccagtcttaagatctttcctgacctgcacc |
| TDP-43 K136TAG forward | cttatggtgcaggtctagaaagatcttaagact |
| TDP-43 K136TAG reverse | agtcttaagatctttctagacctgcaccataag |
| TDP43 K145Q forward | gactggtcattcacaggggtttggctttg |
| TDP43 K145Q reverse | caaagccaaacccctgtgaatgaccagtc |
| TDP43 K145R forward | gactggtcattcaagggggtttggctttg |
| TDP43 K145R reverse | caaagccaaacccccttgaatgaccagtc |
| TDP-43 K79Q forward | gtcaactatccacaagataacaaaagaaaaatg |
| TDP-43 K79Q reverse | catttttcttttgttatcttgtggatagttgac |
| TDP-43 K79R forward | gtcaactatccaagagataacaaaagaaaaatg |
| TDP-43 K79R reverse | catttttcttttgttatctcttggatagttgac |
| TDP43 K84Q forward | gataacaaaagacaaatggatgagacag |
| TDP43 K84Q reverse | ctgtctcatccatttgtcttttgttatc |
| TDP-43 K84R forward | ccaaaagataacaaaagaagaatggatgagacag |
| TDP-43 K84R reverse | ctgtctcatccattcttcttttgttatcttttgg |
| TDP-43 K84TAG forward | ccaaaagataacaaaagatagatggatgagacagat |
| TDP-43 K84TAG reverse | atctgtctcatccatctatcttttgttatcttttgg |
| TDP-43 NheI forward | ctactctagagctagcatgtctgaatatatt |
| TDP-43 NotI reverse | ccccgcggccgcctacattccccagccagaag |
| TDP-43 SalI forward | ggggtcgacgatgtctgaatatattcgggtaaccg |
